# Genomic insights into bacterial isolates dominating honeypot ant crop microbiomes reveal metabolically distinct *Fructilactobacillus* sp.

**DOI:** 10.64898/2026.04.09.717501

**Authors:** Ian M. Oiler, Pranas Grigaitis, Faith A. Hill, Charlotte B. Francoeur, Adria C. LeBoeuf, Lily Khadempour

## Abstract

2.

Honeypot ants engage in a convergently-evolved phenotype called repletism, where specialized workers expand their crops and gasters to store vast amounts of food internally. They then store that food for months to support colonies during times of food scarcity. This fascinating phenotype is not well-understood and only recently have the microbial interactions been studied within the fructose-rich replete crops of these ants. Previous research using amplicon sequencing showed that *Fructilactobacillus* dominates the crop microbiomes of *Myrmecocystus mexicanus* repletes. This striking result and successful isolation of those strains led to the present investigation into the phylogenetic diversity of these strains and any clues to the nature of the symbiotic relationship between them and the ant host. We find that the isolates from these repletes represented two evolutionary lineages, both most closely related to *F*.

*fructivorans*. One of those lineages was also found to be phylogenetically and metabolically distinct from all other *Fructilactobacillus* reference genomes used in this study. We also find evidence for three hypothesized scenarios that could lead to the dominance of these strains in this system, 1. Persistence – the ability of these organisms to exist where others would struggle, 2. Active Competition – additionally producing antimicrobial compounds to reduce competition, and 3. Host Selection – in which the microbe and host share evolutionary history and potential mutualism. These discoveries in a genus of bacteria that are highly relevant for fermented human foods will also lay the groundwork for future understanding of the convergent evolutionary mechanisms of repletism in ants.

**Impact statement:** These analyses add to the literature by identifying a new microbe within a genus that is relevant to food systems. In addition, the host phenotype is convergently evolved and likely microbe-mediated (or at least highly exposed). Understanding this system allows for the testing of ideas of coevolutionary hypotheses with natural replicates. We expect interest to come from food safety and probiotics researchers, evolutionary biologists that think about the impacts of microbes, microbial ecologists interested in novel systems, and those interested in bacteria that may display unique metabolic possibilities. This output allows for clear future examination of this system with many testable hypotheses. Our analysis allows for the creation of a new and unique model system of host-microbe symbiosis.

**Data summary:** New genomes assembled in this work can be found under BioProject ID PRJNA1449409. https://www.ncbi.nlm.nih.gov/bioproject/?term=PRJNA1449409. Reference bacterial genomes were obtained from NCBI at the following accession numbers: SAMN20557570, SAMN28081009, SAMN28081010, SAMN02597458, SAMN43111088, SAMN28081011, SAMN28535231, SAMN04505734, SAMN28081013, SAMN33452149, SAMN37926504, SAMN02849426, SAMN02470196, SAMN02369432, SAMN02797779, SAMN02797782, SAMN02797768, SAMN09762388, SAMN12785275, SAMEA117660288.

**Repositories:** New genomes produced in this study are deposited at the BioProject ID PRJNA1449409.

## 5. Introduction

More than 15,000 species of ants have evolved and adapted to ecosystems on all continents except Antarctica, and they have been diversifying since at least the cretaceous^1,2^. As eusocial insects, ants use division of labor with different colony members, or castes, taking on specialized tasks. This allows members of colonies to efficiently cooperate to both physiologically and behaviorally adapt to a variety of niches. Ants’ evolutionary innovations have allowed them to exploit nearly every biome on Earth^3^. Some fascinating innovations include those of fungus-farming ants, that started farming 65 million years ago^4^, ants whose gut bacteria recycle nitrogen to make essential amino acids^5^, and ants that engage in plant defense and reside inside plant domatia^6^. Our focus is on honeypot ants, which appear highly adapted to fluctuations in seasonal food availability through the repletism phenotype. In this adaptation, specialized ants called repletes are able to store food in excess by expanding their crop many times its normal size (Figure 1). In some cases, repletes become sedentary and are fed mouth-to-mouth via trophallaxis by workers, which forage sugary plant fluids such as nectar and hemipteran honeydew.

**Figure 1.**
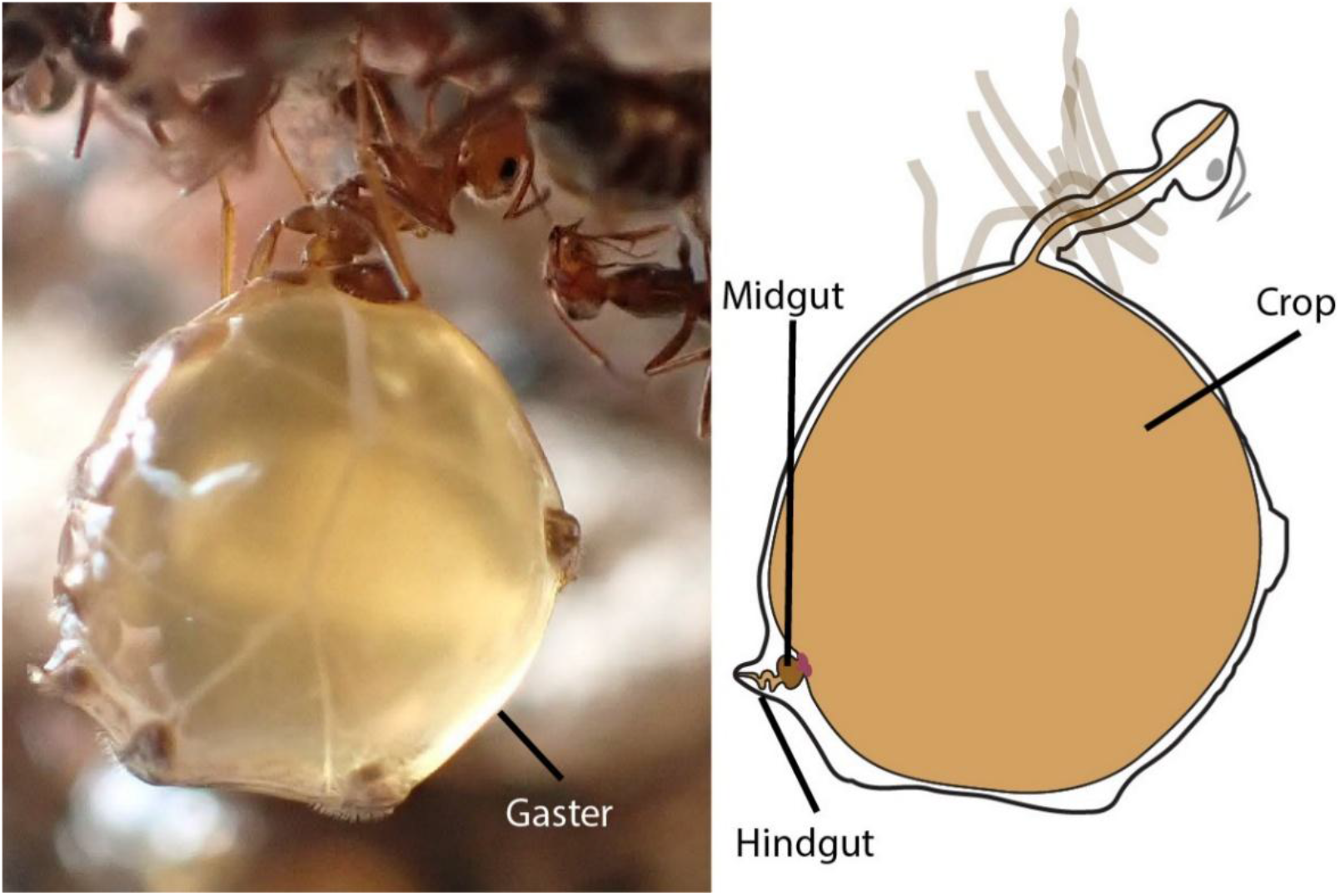
*Myrmecocystus* repletes have specialized morphology for long-term fluid storage in their crops. Left: a photograph of a *M. mendax* replete where the enlarged gaster is clearly visible (photo: M.P. Meurville). Right: Schematic of replete showing the enlarged crop, and relatively small midgut and hindgut.

The repletism phenotype is convergent across at least eight independent ant lineages^7^. Convergent traits provide opportunities to understand instances where evolutionary processes may be predictable. In the case of honeypot ants, there is a clear adaptation to food availability via long-term storage. Honeypot repletes regurgitate their crop contents during times of scarcity to feed the rest of the colony^7^. It is hypothesized that there is a relationship between resource scarcity and repletism. However, there is another layer to this convergently-evolved trait. The sugary liquid food is likely to attract microbes that might lead to spoilage over the many months that these repletes can maintain their stores. Relationships with bacteria and fungi are well documented in some ant lineages (i.e. *Cephalotes^5^, Atta*^4^, *Azteca*^8^). In all animals and plants, microbe-host relationships are ubiquitous and range from pathogens to mutualists, and can be obligate to transient^9,10^. Due to the convergent nature of repletism, it is possible that there are taxonomic or functional similarities in the microbes associated with the crop contents of different honeypot ant species.

Examination of the microbiomes associated with honeypot ants has only been explicitly collected in two lineages. First, in *Camponotus inflatus* in Australia^11^, and second in *Myrmecocystus* spp. in the United States^12^. In *C. inflatus* the whole bodies of only seven ants were used, which resulted in almost exclusively finding the known *Blochmannia* endosymbiont. Of the remaining bacteria (all less than 1% relative abundance) there was a significant proportion of *Apilactobacillus*^11^, a genus of lactic acid bacteria often found in bee samples.

More extensive sequencing of bacterial communities^12^ in several species of *Myrmecocystus* over multiple years, showed *Fructilactobacillus* and other lactic acid bacteria as common gut inhabitants. One species of ant, *M. mexicanus*, was collected in four different sampling years, across three US states, and eight colonies, there was consistently high *Fructilactobacillus* with an average relative abundance of 88.51%. There are many *Fructilactobacillus* species that are common in human food preservation, processing, and spoilage^13–16^. Inclusion in this group of lactic acid bacteria alone means that these microbes are likely of interest to humans.

Additionally, ants in the subfamily *Formicinae* have historically been used in milk fermentation for yogurt in Turkey and Bulgaria due to their acidity and gut resident lactic acid bacteria^15^. Research into honeypot ants allows researchers to test theories of convergent evolution, understand associated microbes that are known to be important to human health and commerce, and represents a tractable framework for understanding the precise mechanisms of host-microbe symbioses.

In the present study, we investigate the *Fructilactobacillus* bacteria from the honeypot ant, *M. mexicanus*, using a genomic approach on isolates from the crop fluid. We wanted to better understand the *Fructilactobacillus* strains that dominate these ant crops, to lay the groundwork for future mechanistic understanding of the host-microbe symbiosis. Through our analysis, we narrowed our hypotheses regarding the mechanism of microbial success in these environments to the following mutually non-exclusive three. 1. Persistence: *Fructilactobacillus* succeeds in a harsh environment where other available microbes cannot. 2. Active Competition: the microbes produce metabolites, like lactic acid, ethanol or polyketides, that further act as a barrier to competition. 3. Host Selection: these microbes provide something through fermentation, or another process, that has led to the host ant either behaviorally or physiologically selecting for this microbe’s success. To do this we provide a thorough description of the genomes and phylogenetic placement of these isolates and establish a predictive framework of microbial functions for future study in this unique system.

## 6. Methods

### Sample Collection

Samples were collected in July 2022 and July 2023 in and around Portal, Arizona (Table 1), either on private property (with permission) or on public Bureau of Land Management (BLM) land. Colonies were found opportunistically and the species of the ants were determined onsite, and later confirmed through COI gene sequencing. Colonies were hand-dug to obtain live repletes. These were delicately removed using forceps and/or paintbrushes into containers. We kept the repletes alive until dissection. At that time, the ants’ heads were removed, and the crop was dissected under sterile conditions. To do this, a 70% ethanol rinse was used to clean the bodies, and the gaster was opened using sterile ultra-fine forceps (Dumont #5SF, FST 11252-00) without penetrating the interior contents of the gaster. Then the entire digestive tract was removed from the gaster and on a sterile petri dish with sterile phosphate buffered saline (PBS). With newly sterilized forceps, the crop was separated and placed in a sterile tube with 100 μL of PBS buffer. The crop was then pierced in the tube using a sterile pipet tip. This resulted in 120 μL to 500 μL of material, which was separated into four new vials; 1) 10 μL for metabolomics, 2) 10 μL with protease inhibitor for proteomics, 3) 10 μL into 90 μL of PBS buffer for microbial growth and isolation, and 4) the rest was reserved for DNA extraction for community amplicon sequencing. For more details on sample collection and dissections, see Nguyen et al.^12^.

**Table 1:** Demographic Information for each sample used in this study.

| Species | Strain | Source | Location | Coordinates | Year | BioSample Acc. |
| --- | --- | --- | --- | --- | --- | --- |
| *A. kunkeei* | DSMZ 12361 | - | USA | - | - | SAMN20557570 |
| *F. carniphilus* | KI4_A6 | insect | Kangaroo Island, AUS | -35.83305 , 137.564317 | 2019 | SAMN28081009 |
| *F. cliffordii* | K14_B1 | insect | Kangaroo Island, AUS | -35.83305 , 137.564317 | 2019 | SAMN28081010 |
| *F. florum* | 8D | fruit | California, USA | - | - | SAMN02597458 |
| *F. frigidiflavus* | FUA3702 | sourdough | British Colombia, CAN | 49.9142 , -116.9155 | 2019 | SAMN43111088 |
| *F. hinevesii* | KI11_C11 | plant | Kangaroo Island, AUS | -35.852272 , 137.720672 | 2019 | SAMN28081011 |
| *F. ixorae* | Ru20-1 | plant | Kalasin, THA | 16.4314 , 103.5059 | 2019 | SAMN28535231 |
| *F. linderi* | TMW 1.1993 | alcohol | NLD | - | 1993 | SAMN04505734 |
| *F. myrtifloralis* | KI16_H9 | plant | Kangaroo Island, AUS | -35.745378 , 136.876892 | 2019 | SAMN28081013 |
| *F. sanfranciscensis* | SPC-SNU-70-4 | alcohol | Seoul, KOR | - | 2021 | SAMN33452149 |
| *F. vespulae* | Mu01 | plant | Valencia, ESP | - | 2020 | SAMN37926504 |
| *F. fructivorans* | DmCS_002 | insect | New York, USA | - | 2011 | SAMN02849426 |
| *F. fructivorans* | KCTC 3543 | - | - | - | - | SAMN02470196 |
| *F. fructivorans* | DSM 20203 | - | - | - | 1934 | SAMN02369432 |
| *F. fructivorans* | DSM 20350 | food | - | - | 1934 | SAMN02797779 |
| *F. fructivorans* | DSM 20571 | alcohol | JPN | - | 1957 | SAMN02797782 |
| *F. fructivorans* | ATCC 27394 | food | - | - | 1934 | SAMN02797768 |
| *F. fructivorans* | KCTC 3543 (2) | - | - | - | - | SAMN09762388 |
| *F. fructivorans* | LF543 | - | - | - | 1990 | SAMN12785275 |
| *F. fructivorans* | CIP103141 | - | - | - | - | SAMEA117660288 |
| *F. sp1* | X1R3C1 | insect | Arizona, USA | 31.908 , -109.152 | 2022 | SAMN57118716 |
| *F. sp1* | X1R3C2 | insect | Arizona, USA | 31.908 , -109.152 | 2022 | SAMN57118717 |
| *F. sp1* | X2R1C1 | insect | Arizona, USA | 31.902 , -109.109 | 2022 | SAMN57118718 |
| *F. sp1* | X2R1C3 | insect | Arizona, USA | 31.902 , -109.109 | 2022 | SAMN57118719 |
| *F. sp2\** | X2R5C2 | insect | Arizona, USA | 31.902 , -109.109 | 2022 | SAMN57118720 |
| *F. sp1* | X2R5C3 | insect | Arizona, USA | 31.902 , -109.109 | 2022 | SAMN57118721 |
| *F. sp1* | X2R6C1 | insect | Arizona, USA | 31.902 , -109.109 | 2022 | SAMN57118722 |
| *F. sp1* | X3R2C1 | insect | Arizona, USA | 31.902 , -109.109 | 2022 | SAMN57118723 |
| *F. sp1* | X3R2C3 | insect | Arizona, USA | 31.902 , -109.109 | 2022 | SAMN57118724 |
| *F. sp1* | X3R5C2 | insect | Arizona, USA | 31.902 , -109.109 | 2022 | SAMN57118725 |
| *F. sp1* | X7R4C1 | insect | Arizona, USA | 32.0356333 , -109.1945333 | 2023 | SAMN57118726 |
| *F. sp1* | X7R5C1 | insect | Arizona, USA | 32.0356333 , -109.1945333 | 2023 | SAMN57118727 |
| *F. sp1* | X7R7C1 | insect | Arizona, USA | 32.0356333 , -109.1945333 | 2023 | SAMN57118728 |
| *F. sp1* | X7R9C1 | insect | Arizona, USA | 32.0356333 , -109.1945333 | 2023 | SAMN57118729 |
| *F. sp1* | X8R1C1 | insect | Arizona, USA | 32.035083 , -109.193433 | 2023 | SAMN57118730 |
| *F. sp1* | X8R2C1 | insect | Arizona, USA | 32.035083 , -109.193433 | 2023 | SAMN57118731 |
| *F. sp1* | X8R4C1 | insect | Arizona, USA | 32.035083 , -109.193433 | 2023 | SAMN57118732 |

\*isolate is not monophyletic with the rest of the isolates and is likely *F. fructivorans* (Figure 2 for more)

### Sample Preparation, Sequencing, and Genome Curation

Upon initial analysis of community amplicon sequencing data, we observed that *Fructilactobacillus* dominated the crops of *M. mexicanus* ants. Therefore, strains of this genus were isolated. Initially, through 16S rRNA gene sequencing, 17 unique strains were identified across 13 different repletes, from five colonies (X1, X2, X3, X7, X8) and their genomes were sequenced. *Fructilactobacillus* colonies were isolated from the material in the PBS buffer. Pure isolates were grown overnight in De Man, Rogosa, and Sharpe (MRS) broth and then centrifuged to form a pellet and then stored at −20°C until DNA extraction. DNA extraction was done using the DNeasy Blood and Tissue Kit (QIAGEN, Aarhus, Denmark) with the default instructions for the Gram-positive pretreatment (30 minute incubation time). Samples were sent to SeqCoast Genomics (Portsmouth, NH, USA) for whole genome sequencing, where samples were prepared using the Illumina DNA Prep tagmentation kit using unique dual indexes. Sequencing was performed on the Illumina NextSeq2000 platform using a 300-cycle flow cell kit to produce 2×150bp paired reads. 1-2% PhiX control was spiked into the run to support optimal base calling. Read demultiplexing, read trimming, and run analytics were performed using DRAGEN v3.10.12, an on-board analysis software on the NextSeq2000.

Reads were processed using FASTP^17^ and assembled with SPAdes v3.15.2^18^. We assessed the completeness and contamination levels of the genomes using BUSCO^19^ and checkM^20^ using the Lactobacillus (UID436) marker lineage. Coverage was calculated with BBMap^21^. We removed contigs with less than 200bp. We retrieved nine *Fructilactobacillus fructivorans*, 10 *Fructilactobacillus* species (*F. carniphilus, F. cliffordii, F. hinevesii, F. myrtifloralis, F. ixorae, F. florum, F. sanfranciscensis, F. frigidiflavus, F. vespulae, and F. linderi*), and one *Apilactobacillus kunkeei* (outgroup) genome assemblies from the NCBI RefSeq database to compare to our isolated genomes (Table 1 for more information). These represent all of the reference quality genomes on NCBI for *Fructilactobacillus* (at the time of retrieval) as well as eight non-reference quality genomes for *F. fructivorans* to inspect the closest relative further.

### Comparative Genomics and Modeling

Comparative genomics analyses were done using Anvi’o v9^22^, beginning with creating a genome database for all 37 genomes using ‘anvi-gen-genomes-storage’. This includes gene calling done by Prodigal^23^ to identify COG20 annotations^24^. Genomes were then compared using the ‘anvi-pan-genome’ command using default settings except MCL inflation was set to 10 (for closely related genomes). This command entailed identifying similarities between amino acid sequences using DIAMOND^25^, performing multiple sequence alignments of the amino acid sequences in each orthogroup, computing functional and geometric homogeneity, computing average amino acid identity within each orthogroup, and organizing the orthogroups using hierarchical clustering based on their distribution across genomes. Enriched gene functions were calculated in certain groups of genomes using the ‘anvi-compute-functional-enrichment-in-pan’ command. Genomes were separated into groups for functional enrichment based on the labeling of the phylogeny (Figure 2). The Bacteria_71 concatenated gene set in Anvi’o was used to construct a phylogeny using IQtree2^26^. We calculated genome similarity by ANI score using ‘anvi-compute-genome-similarity’. The command ‘anvi-estimate-metabolism’ was used to quantify the stepwise completeness of metabolic pathways using the KEGG database^27^.

**Figure 2.**
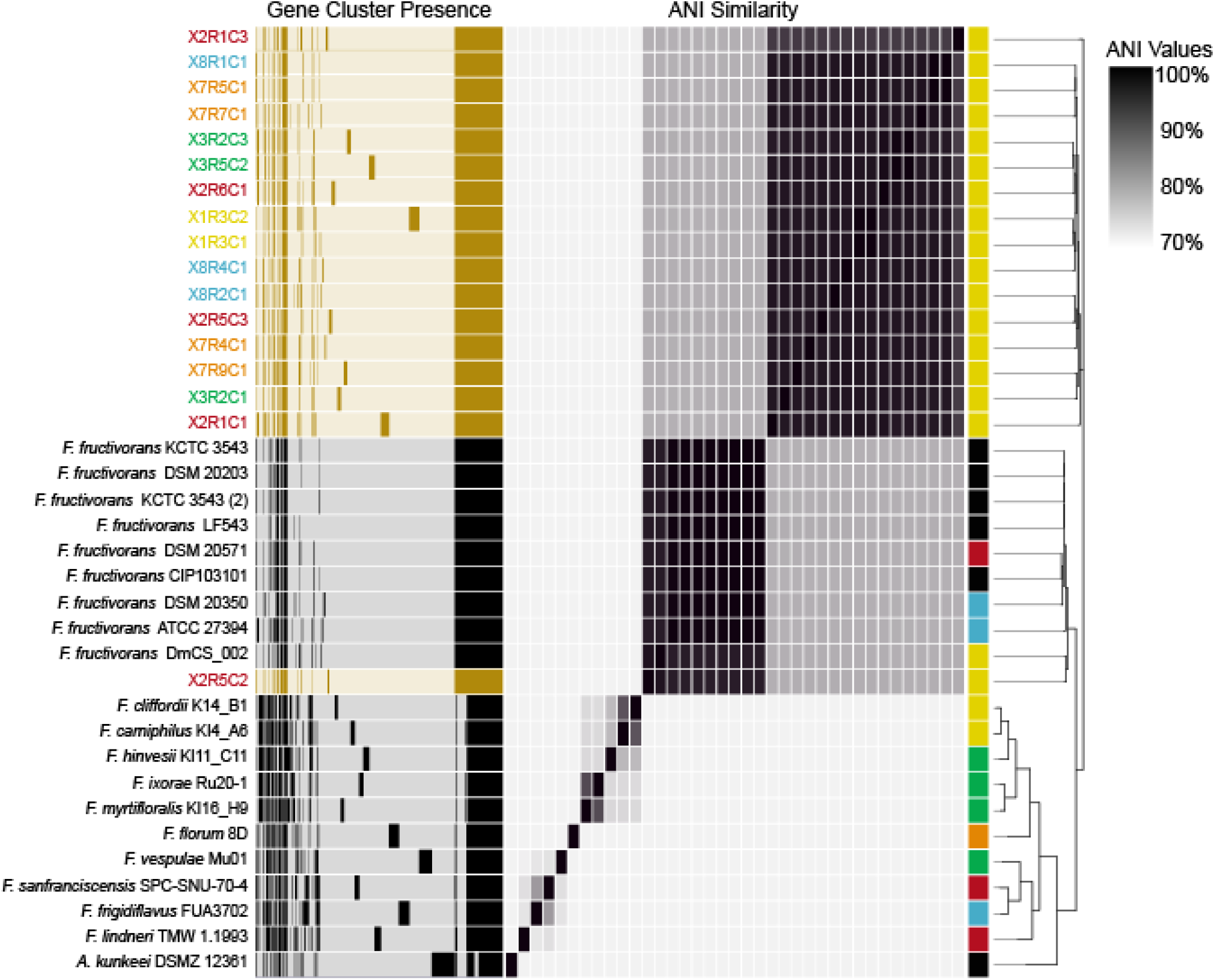
Isolated *Fructilactobacillus* samples fall into two distinct lineages, most closely related to *F. fructivorans*. From left to right, Sample names with isolates colored by colony, Gene presence/absence clustering, ANI similarity heatmap (values in text and Supplement 2), sample source (orange = fruit, yellow = insect, green = plant, red = alcoholic beverage, blue = food, black = unknown), and genomic phylogeny used to order samples.

**Figure 3.**
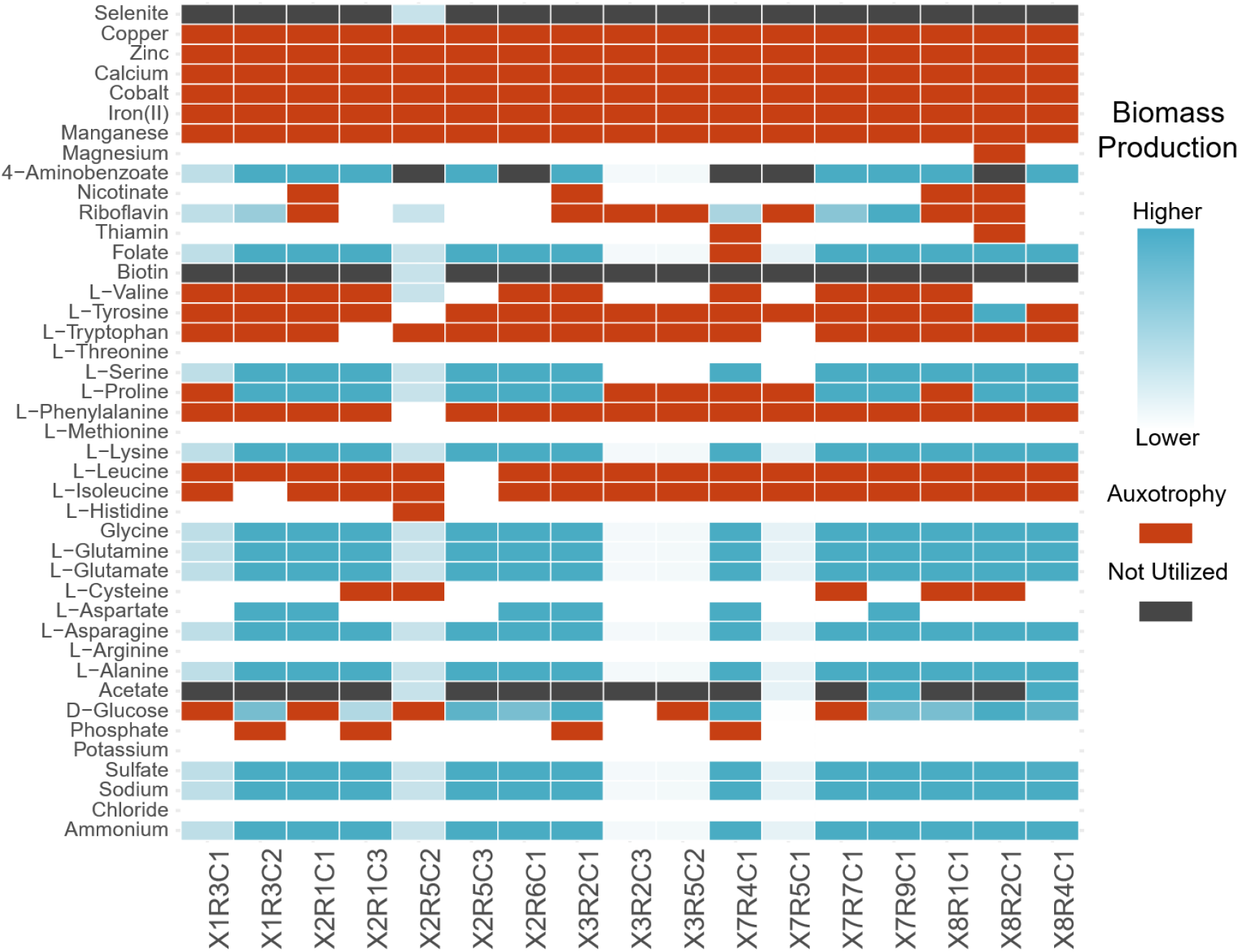
Measurement of predicted biomass production under resource scarcity shows multiple shared auxotrophies across all isolate metabolic models. Blue gradient indicates biomass production when each metabolite’s uptake is turned off. Red indicates no biomass produced (metabolite is necessary – auxotrophy). Black indicates that the model does not use that metabolite for biomass production. The gradient goes from low production (white) to higher production (blue).

We also created genome-scale metabolic models (GEMs) to further compare the metabolic potential of sequenced *Fructilactobacillus* isolates. Genomes were not processed with the Anvi’o pipeline described above. Open reading frames (ORFs) were called from the genome sequences and functionally annotated using Bakta^28^. We used CarveMe^29^ to create draft genome-scale metabolic models from the protein FASTA files using the default settings, except for specifying the Gram-positive universe and using the SCIP SoPlex linear program solver^30^. The draft models were gapfilled in CarveMe on chemically defined MRS media (Supplementary Data 1), with which growth has been confirmed for each isolate strain. The GEMs were then imported into COBRApy^31^ in Python version 3.10 for comparisons of model metrics (metabolites, reactions, metabolic genes) and predicting growth auxotrophies.

We used the tool, antiSMASH 8.0^32^ to search our genomes for biosynthetic gene clusters that may produce secondary metabolites. For each submitted genome, default parameters for bacteria were used.

## 7. Results and Discussion

The goal of this work is to generate and clarify hypotheses about the functional relationship between the *M. mexicanus* host and *Fructilactobacillus* microbial isolates. Our interest was in better understanding the nature of this symbiosis, spurred by the findings that these microbes dominate the crop microbiomes of this host species over dozens of samples, multiple colonies and four sampling years. It is therefore unlikely that this pattern is the result of a fully stochastic process and instead is likely due to strong mechanisms driving this pattern. Notably, close relatives of *M. mexicanus*, living sympatrically with them and presenting repletism, do not show such a strong pattern of dominance for any microbe, though the genus *Fructilactobacillus* and other lactic acid bacteria are common among those communities as well (using 16S rRNA gene sequencing^12^).

The nature of this symbiosis is unknown (parasitic, mutualistic, commensal, etc.). To assess this, we assembled the genomes of 17 isolates from 13 honeypot crop samples, spanning five different colonies, and compared them to the genomes of 20 close bacterial relatives. This was done with the goal of better classifying and predicting the functional capabilities of these isolates. The range of genome lengths were 1,334,748 - 1,487,218bp, with an average size of 1,392,421 and average coverage of 305X. The average %GC was 38.95%. BUSCO completeness of all isolates was over 98.8% and checkM completeness was 99.37% for all genomes. By comparing these genomes’ phylogenetic relatedness, broad gene content, and draft metabolic models, we begin to assess the likelihood of our three hypothesized conditions (1. Persistence, 2. Active Competition, and 3. Host Selection). In fact, we find evidence for all three conditions and discuss below when that evidence provides support for each of these hypotheses.

### Phylogenetic Placement

Phylogenetic analysis shows that one isolate is *Fructilactobacillus fructivorans* and the rest are in a separate but sister lineage (Figure 2). The isolate strain, X2R5C2, was found to be sister to the reference assembly for *F. fructivorans* isolated from the intestinal tract of *Drosophila melanogaster* instead of being monophyletic with the rest of the isolates. Average Nucleotide Identity (ANI) was calculated for each genome, and confirms the pattern seen in the phylogeny (Figure 2). Specifically, X2R5C2 is 98% identical to the *F. fructivorans* reference genome (DmCS_002 in Figure 2) and between 80% and 83% identical to the rest of the isolate lineages. Using a 95% threshold for putative species^33^, another isolate, X2R1C3, barely misses this cut, but is still monophyletic with the rest of the honeypot clade (though on the outer edge). The rest of the isolates are 97-99% similar by ANI to each other, 81-82% similar to the *F. fructivorans* clade and 70-73% similar to the rest of the tested genomes, including *A. kunkeei* (see Supplement 2 for ANI matrix).

The results of the phylogenetic analysis provide some initial clues as to what environments these microbes are likely to thrive in as well as the possible functions that they may serve in those environments. We already knew that these isolates fell within the genus *Fructilactobacillus* from previous amplicon sequencing^12^. However, their closest relatives being *F. fructivorans* already presents evidence for the Persistence hypothesis as these microbes are known to survive harsh conditions of low pH and high ethanol content^34^. In addition, they are heterofermentative lactic acid bacteria that often quickly increase the acidity in their environments, reducing competition, thereby and supporting the Active Competition hypothesis. Host Selection might also explain why most of these strains could be considered part of the same population by ANI score, despite coming from five different colonies of ants across multiple locations. While much can be inferred from the phylogenetic placement of these microbes, we also used annotation tools to directly predict functional traits.

In comparing shared orthogroups between genomes and their likely functional associations, we found 583 orthogroups that all genomes in the dataset share. There were 137 additional orthogroups shared among the *Fructilactobacillus* genus (only excluding *A. kunkeei*) and 213 orthogroups shared only between the *F. fructivorans* and the honeypot clade. Finally, there were 22 orthogroups that were shared only within the monophyletic clade of honeypot isolates (which excludes X2R5C2).

We performed functional enrichment tests to identify orthogroup functions that were more commonly found in some genomes over others. There were multiple functions that were unique to different groups. For the honeypot clade (excluding X2R5C2), this included only two orthogroup functions, Zn-dependent alcohol/formaldehyde dehydrogenase (FrmA) and DNA-binding transcriptional regulator YiaG, XRE-type HTH domain (YiaG). There were no functions in the *F. fructivorans* clade (including X2R5C2) that were not shared with at least one other *Fructilactobacillus* genome. For the rest of the genomes, there were 7 functions that were found in all genomes except the *F. fructivorans* and honeypot clades, representing potential losses of function. These were 3-keto-L-gulonate-6-phosphate decarboxylase (UlaD), AAA+-type ATPase, SpoVK/Ycf46/Vps4 family (SpoVK), Adenosine deaminase (Add), D-arabinose 5-phosphate isomerase (GutQ), 1,4-Dihydroxy-2-naphthoyl-CoA synthase (MenB), Phytoene/squalene synthetase (ERG9), Acyl-CoA thioesterase (PaaI), and two uncharacterized proteins. We found a total of 322 enriched functions for various groups of genomes, which can be found in Supplementary Data 3.

The FrmA orthogroup is unique to the honeypot ant clade. This is a Zn-dependent alcohol/formaldehyde dehydrogenase, which can catalyze the reaction between ethanol and formaldehyde, or between formic acid and formaldehyde^35^. Notably, the ants, which belong to the formicine subfamily, produce and consume formic acid, which is related to controlling their gut microbial communities^36^. The honeypot ant clade-specific presence of this orthogroup could be an adaptation to the formicine ant gut environment. There are other orthogroups that are found in all honeypot ant strains but are not unique to them: acid resistance membrane protein (HdeD), NAD dependent aldehyde dehydrogenase (AdhE), and antitoxin component YafN of the YafNO toxin-antitoxin module (StbD). These could be important if the microbes serve a function in detoxifying plant defense compounds from the plant fluid food sources. However, not only are some of these annotations unspecific, we also cannot conclude that these functions are occurring without experimentation. In general, more work needs to be done to understand how these gene functions interact in the honeypot crop environment, but there appears to be some metabolic differences between the honeypot isolates and other members of *Fructilactobacillus*.

### Metabolic Modeling

To quantify the metabolic potential of *Fructolactibacillus* isolates, we reconstructed genome scale metabolic models (GEMs) using the automated pipeline CarveMe. We performed automatic gap filling of the GEMs for growth on MRS medium, after confirming growth for all isolates on this chemically-defined media. The demographic values (genes, reactions, and metabolites) were somewhat variable between isolates. The number of genes ranged from 357 to 384, metabolites ranged from 642 to 788, and reactions ranged from 952 to 1133 (detailed demographic information in Supplementary data 4). These demographic values were not obviously associated with the colony of origin or patterns found in the phylogeny. The GEMs were used to identify predicted auxotrophies, or molecules needed for cell growth, by iteratively shutting off uptake for each molecule from our in-silico MRS media and measuring flux with COBRApy using the equation for biomass. Our analysis found that most molecules were able to be produced at some level in the honeypot isolates. However, there are multiple metals found to be auxotrophies across all or most honeypot samples, as well as the following amino acids; tyrosine, tryptophan, phenylalanine, leucine, and isoleucine. Future analyses of host contributions and detailed identification of the honeypot liquid chemistry will allow for better understanding about where these auxotrophies are being supplied. In the case of the Host Selection hypothesis, we would expect that some of these auxotrophies are produced and secreted into the crop by the host ants or through their dietary intake.

We estimated the stepwise completeness of various metabolic pathways across all samples using the KOfam database (Figure 4). A pathway was considered complete if it contained 75% of the metabolic steps from known KEGG references. Thirty-nine pathways were complete in at least one sample and 14 were considered complete in all samples. Of the other 25, our honeypot isolates tended to cluster with the *F. fructivorans* group and in one instance, X2R5C2 (the only genome that falls within the *F. fructivorans* clade) shared a ketone body biosynthesis pathway that was only found in some *F. fructivorans* samples. Figure 4 shows that many pathways are conserved between all, or almost all, genomes across the dataset. However, there are many strain-specific pathways not found in the honeypot ant clade, throughout the phylogeny. These pathways could be specific adaptations to the environment that these strains are found in.

**Figure 4:**
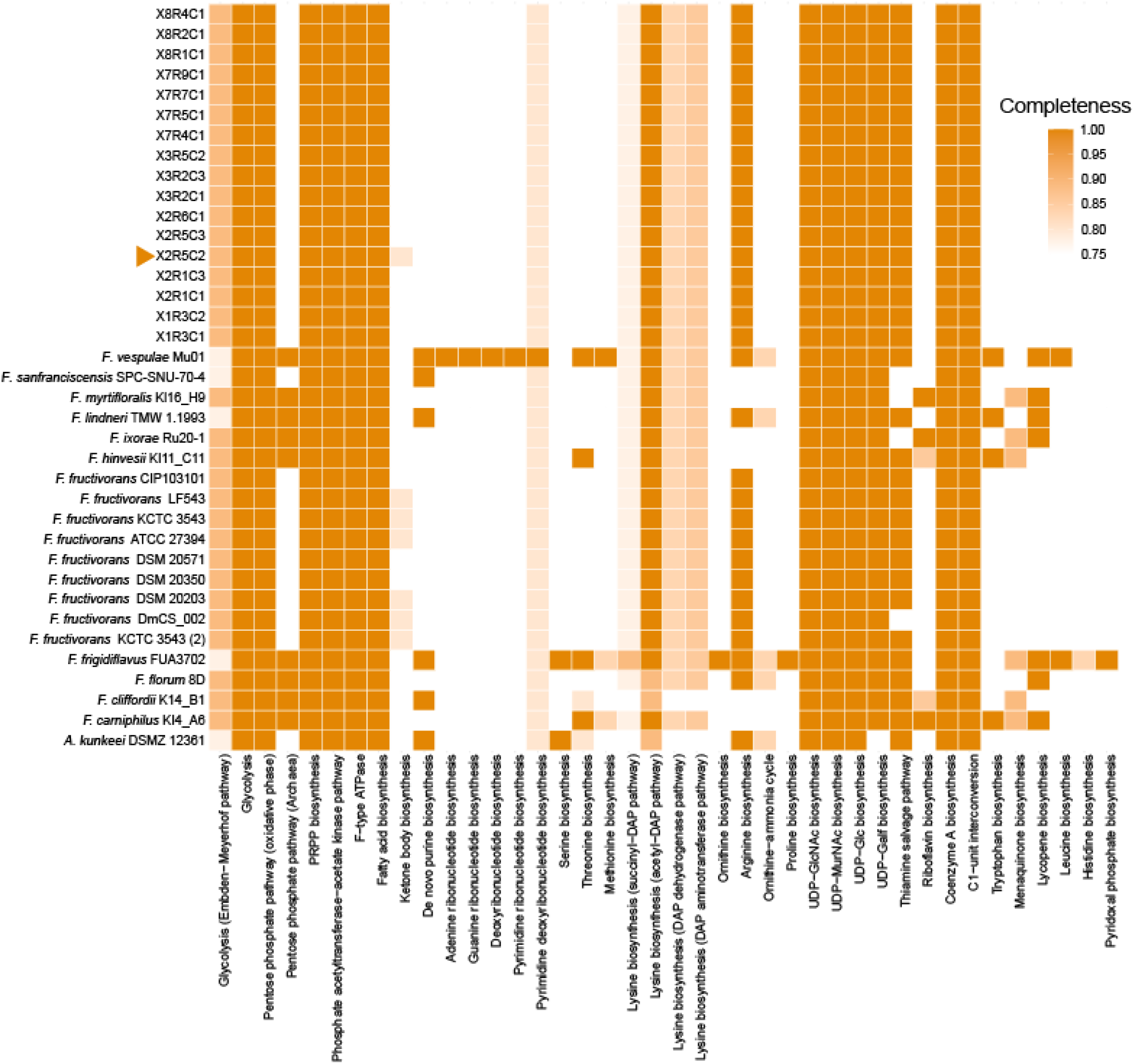
Stepwise completeness of KEGG modules for each genome shows many shared complete pathways, with some clustering within phylogenetically distinct groups. Completeness was defined as the proportion of pathway steps there is evidence for in the genomes. For the purpose of this study, pathways with more than 75% completeness were considered to be complete. X2R5C2 is marked with an arrow as it showed distinct differences in other analyses and shares a complete pathway in this analysis that exclusively shows up in *F. fructivorans* (ketone body biosynthesis).

Using antiSMASH, we found three biosynthetic gene clusters (BGCs) that were conserved across the honeypot ant isolate clade. These were a terpene precursor, Type III polyketide synthase (T3PKS), and lanthipeptide class III. All honeypot isolates were positive for terpene precursors and T3PKS. All but X2R5C2 and X2R1C3 were positive for the lanthipeptide class III BGC (Table 2). The terpene precursor and T3PKS BGCs were shared among all genomes. None of the non-honeypot ant *Fructilactobacillus* genomes had the lanthipeptide class III BGC. However, a separate terpene BGC was unique to the *F. fructivorans* group (including X2R5C2) and most of the other *Fructilactobacillus*, but not the monophyletic group of honeypot isolates or *Apilactobacillus kunkeei*. The presence of different compounds that might be used in defense or antimicrobial competition provides evidence for the Active Competition hypothesis. If the bacteria are actively producing these compounds, they might be acting as additional deterrents for the growth of competitive microbes, one addition to deterrence through lactic acid production. This is a hypothesis that can be verified with future metabolomic analyses of the honeypot crop fluid and lab strain testing.

**Table 2:**
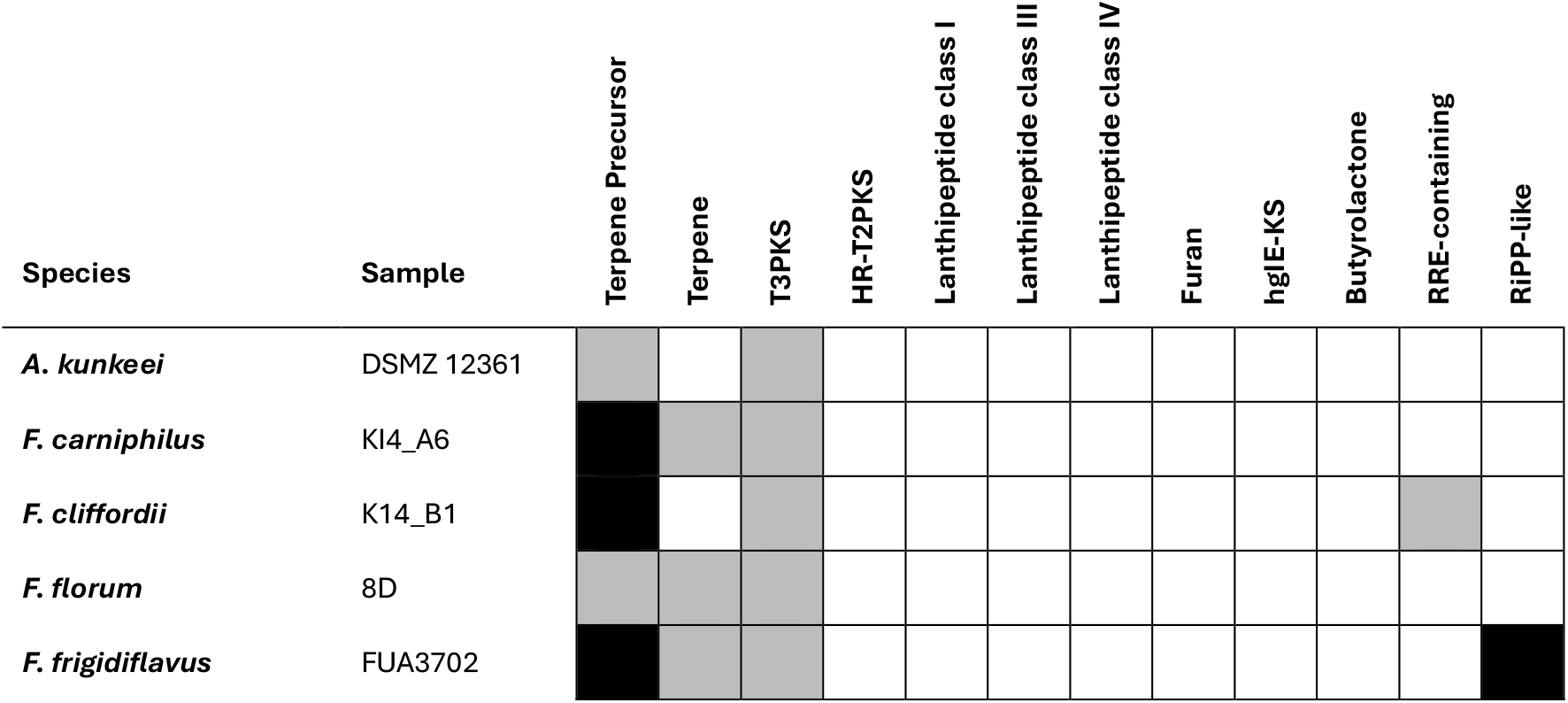
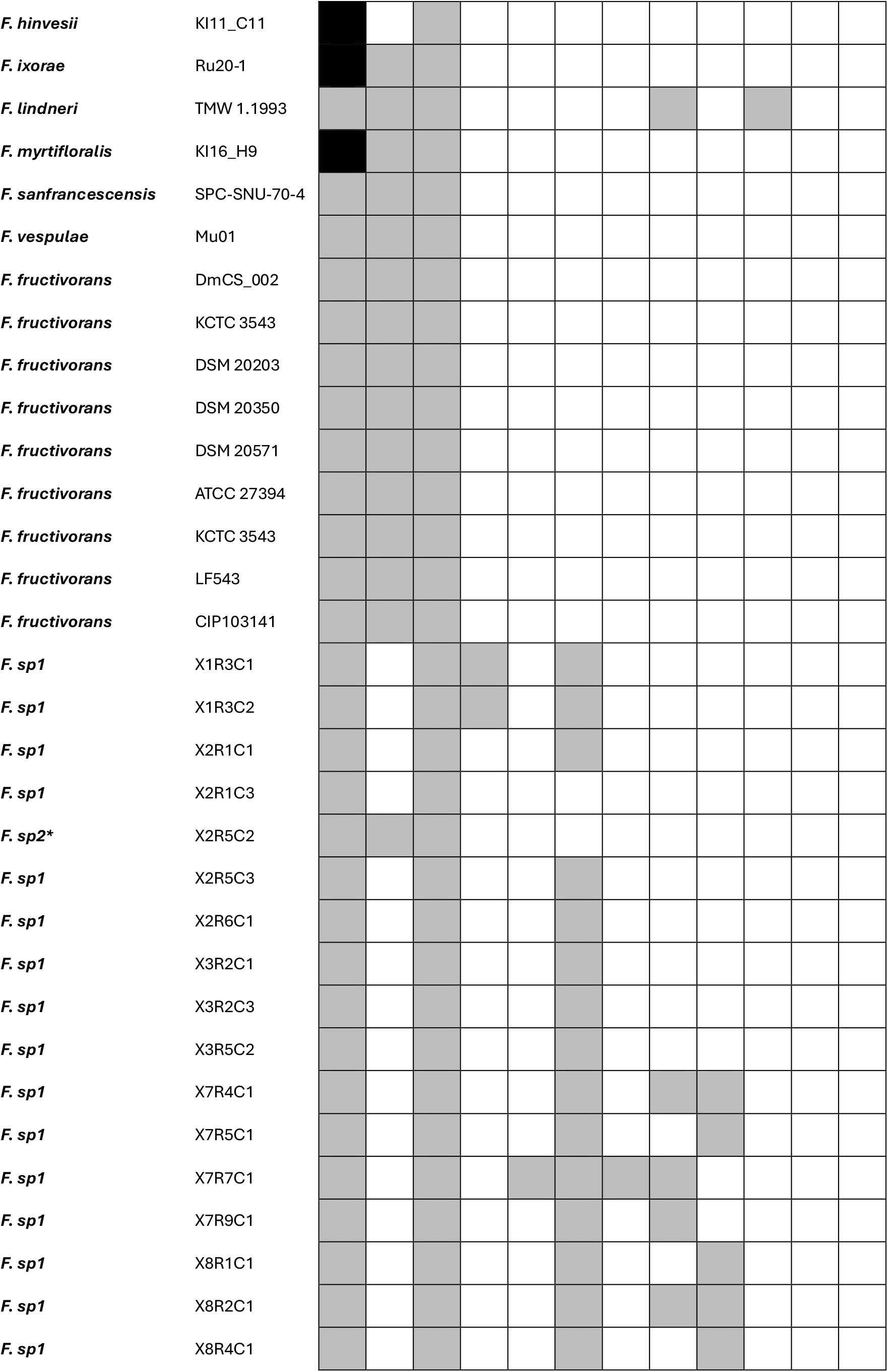
AntiSMASH biosynthetic gene clusters confirm differences between phylogenetic groups based on secondary metabolite production. White squares = no hits for that cluster, gray squares = single hit, black squares = two hits.

## Conclusions

*Fructilactobacillus* contains multiple species that are commonly found in, or explicitly used for, human foods and environments. In every case, these microbes have evolved in nature and have been utilized by humans. Increased knowledge of these microbes will always be beneficial to human health and society, and better understanding their presence in novel environments can only expand that benefit. In this study, we examine the genomes of the honeypot ant symbiont *Fructilactobacillus* and begin the process of predicting their impacts in a novel host-microbe symbiosis. We find that most honeypot isolates represent metabolically distinct lineages compared to the reference genomes analyzed. We find that their predicted functions, from phylogenetic inference, genome comparisons, and metabolic modeling provide evidence for all three hypothesized reasons for their dominance of *M. mexicanus* crop microbiomes. The hardiness of close relatives and the presence of genes for acid tolerance indicate support for simply persisting where other microbes could not. The presence of additional biosynthetic clusters for defense-related compounds and their assumed production of lactic acid to further lower pH indicates evidence for the hypothesis of Active Competition. Finally, the potential loss of some functional pathways, auxotrophies for some important metals and amino acids, and the presence of a unique formaldehyde-detoxifying orthogroup function maybe indicate more specialization for this host niche and suggests possible coevolutionary history. The intention of these analyses was to build a scaffold that can be used in future studies for more precise mechanistic analyses within this specific system, the extended network of convergent repletism, and symbiotic systems more broadly. Our future work will include characterization and development of the honeypot liquid media for in vitro experiments, iteratively improving the genome scale metabolic modeling effort, and lab experiments to confirm the resource needs and metabolic products of the *Fructilactobacillus* isolates.

## Supporting information

Supplement 1 - MRS Media

Supplement 2 - ANI Matrix

Supplement 3 - Enriched Functions

Supplement 4 - GEM Stats

## 8. Author statements

### 8.1 Author contributions

Conceptualization was primarily done by IMO and LK, with help from PG and ACL. Data Curation was done by CF, IMO, FAH, and LK. Formal analysis was done by IMO with help from PG. Funding acquisition was done by LK, CF. Writing was primarily done by IMO with review and editing from LK, PG, ACL.

### 8.4 Ethical approval

Not applicable as we did not conduct experiments with humans or vertebrate animals.

### 8.5 Consent for publication

We did not perform our study on human subjects.

## 8.6 Acknowledgements

This work was completed as part of the Rutgers Artificial Intelligence and Data Science (RAD) Collaboratory and made possible with funding and support provided by the Rutgers-New Brunswick Office of the Chancellor, the Rutgers-New Brunswick Office for Research, and the Rutgers University Office of the Executive Vice President for Academic Affairs acquired by IMO.

## References

1. AntWeb. Accessed March 29, 2026. https://www.antweb.org/

2. Schultheiss P, Nooten SS, Wang R, Wong MKL, Brassard F, Guénard B. The abundance, biomass, and distribution of ants on Earth. Proc Natl Acad Sci U S A. 2022;119(40):e2201550119. doi:10.1073/pnas.2201550119

3. Toro I Del, Ribbons RR, Pelini SL, Forest H. The little things that run the world revisited: a review of ant-mediated ecosystem services and disservices (Hymenoptera: Formicidae). Myrmecol News. 2012;17(10):133–146.

4. Schultz TR, Sosa-Calvo J, Kweskin MP, et al. The coevolution of fungus-ant agriculture. Science (1979). 2024;386(6717):105–109. doi:10.1126/science.adn7179

5. Hu Y, Sanders JG, Łukasik P, et al. Herbivorous turtle ants obtain essential nutrients from a conserved nitrogen-recycling gut microbiome. Nature Communications 2018 9:1. 2018;9(1):964–. doi:10.1038/s41467-018-03357-y

6. Brouat C, McKEY D. Origin of caulinary ant domatia and timing of their onset in plant ontogeny: evolution of a key trait in horizontally transmitted ant-plant symbioses. Biological Journal of the Linnean Society. 2000;71(4):801–819. doi:10.1111/j.1095-8312.2000.tb01292.x

7. Nogueira BR, Leboeuf AC, Barden P, Nguyen V, Khadempour L. Honeypots: a review of repletism across the ants. Myrmecol News. 2026;36:39–57. doi:10.25849/myrmecol.news_036:039

8. Barrajon-Santos V, Nepel M, Hausmann B, Voglmayr H, Woebken D, Mayer VE. Dynamics and drivers of fungal communities in a multipartite ant-plant association. BMC Biology 2024 22:1. 2024;22(1):112–. doi:10.1186/s12915-024-01897-y

9. Hammer TJ, Sanders JG, Fierer N. Not all animals need a microbiome. FEMS Microbiol Lett. 2019;366(10):117. doi:10.1093/femsle/fnz117

10. Khadempour L. A new conceptual framework for host-microbe symbiosis. Published online February 25, 2026. doi:10.32942/X2K95W

11. Dong AZ, Cokcetin N, Carter DA, Fernandes KE. Unique antimicrobial activity in honey from the Australian honeypot ant (Camponotus inflatus). PeerJ. 2023;11:e15645. doi:10.7717/PEERJ.15645/SUPP-1

12. Diemquynh :, Nguyen V, Francoeur CB, et al. Myrmecocystus honeypot ants have species specific resident gut microbiome. bioRxiv. Published online April 8, 2026:2026.04.07.717087. doi:10.64898/2026.04.07.717087

13. Charlton DB; NME; WCH, Charlton DB, Nelson ME, Werkman CH. Physiology of Lactobacillus fructivorans sp. nov. isolated from spoiled salad dressing. Iowa State Coll Jour Sci. 1934;9(1):1–11. Accessed March 7, 2026. https://eurekamag.com/research/025/243/025243004.php

14. Takahashi M, Morikawa K, Akao T. Novel method for predicting the risk of spoilage by lactic acid bacteria during the storage of Japanese sake. Applied Food Research. 2025;5(1):100835. doi:10.1016/J.AFRES.2025.100835

15. Sinotte VM, Ramos-Viana V, Vásquez DP, et al. Making yogurt with the ant holobiont uncovers bacteria, acids, and enzymes for food fermentation. iScience. 2025;28(10):113595. doi:10.1016/j.isci.2025.113595

16. Ibrahim SA, Ayivi RD, Zimmerman T, et al. Lactic Acid Bacteria as Antimicrobial Agents: Food Safety and Microbial Food Spoilage Prevention. Foods 2021, Vol 10, Page 3131. 2021;10(12):3131. doi:10.3390/FOODS10123131

17. Chen S, Zhou Y, Chen Y, Gu J. fastp: an ultra-fast all-in-one FASTQ preprocessor. Bioinformatics. 2018;34(17):i884–i890. doi:10.1093/bioinformatics/bty560

18. Bankevich A, Nurk S, Antipov D, et al. SPAdes: a new genome assembly algorithm and its applications to single-cell sequencing. J Comput Biol. 2012;19(5):455–477. doi:10.1089/cmb.2012.0021

19. Manni M, Berkeley MR, Seppey M, Zdobnov EM. BUSCO: Assessing Genomic Data Quality and Beyond. Curr Protoc. 2021;1(12). doi:10.1002/cpz1.323

20. Parks DH, Imelfort M, Skennerton CT, Hugenholtz P, Tyson GW. CheckM: assessing the quality of microbial genomes recovered from isolates, single cells, and metagenomes. Genome Res. 2015;25(7):1043–1055. doi:10.1101/gr.186072.114

21. Bushnell B. BBMap: A Fast, Accurate, Splice-Aware Aligner. Published online 2014.

22. Eren AM, Kiefl E, Shaiber A, et al. Community-led, integrated, reproducible multi-omics with anvi’o. Nature Microbiology 2020 6:1. 2020;6(1):3–6. doi:10.1038/s41564-020-00834-3

23. Hyatt D, Chen GL, LoCascio PF, Land ML, Larimer FW, Hauser LJ. Prodigal: prokaryotic gene recognition and translation initiation site identification. BMC Bioinformatics. 2010;11. doi:10.1186/1471-2105-11-119

24. Galperin MY, Vera Alvarez R, Karamycheva S, et al. COG database update 2024. Nucleic Acids Res. 2025;53(D1):D356–D363. doi:10.1093/nar/gkae983

25. Buchfink B, Xie C, Huson DH. fast and sensitive protein alignment using diamond. Published online 2014. doi:10.1038/nmeth.3176

26. Minh BQ, Schmidt HA, Chernomor O, et al. IQ-TREE 2: New Models and Efficient Methods for Phylogenetic Inference in the Genomic Era. Mol Biol Evol. 2020;37(5):1530–1534. doi:10.1093/molbev/msaa015

27. Kanehisa M, Furumichi M, Sato Y, Matsuura Y, Ishiguro-Watanabe M. KEGG: biological systems database as a model of the real world. Nucleic Acids Res. 2025;53(D1):D672–D677. doi:10.1093/nar/gkae909

28. Schwengers O, Jelonek L, Dieckmann MA, Beyvers S, Blom J, Goesmann A. Bakta: Rapid and standardized annotation of bacterial genomes via alignment-free sequence identification. Microb Genom. 2021;7(11):000685. doi:10.1099/MGEN.0.000685/CITE/REFWORKS

29. Machado D, Andrejev S, Tramontano M, Patil KR. Fast automated reconstruction of genome-scale metabolic models for microbial species and communities. Nucleic Acids Res. 2018;46(15):7542–7553. doi:10.1093/NAR/GKY537

30. Achterberg T, Berthold T, Koch T, Wolter K. Constraint Integer Programming: A New Approach to Integrate CP and MIP. Lecture Notes in Computer Science (including subseries Lecture Notes in Artificial Intelligence and Lecture Notes in Bioinformatics). 2008;5015 LNCS:6-20. doi:10.1007/978-3-540-68155-7_4

31. Ebrahim A, Lerman JA, Palsson BO, Hyduke DR. COBRApy: COnstraints-Based Reconstruction and Analysis for Python. BMC Systems Biology 2013 7:1. 2013;7(1):74–. doi:10.1186/1752-0509-7-74

32. Blin K, Shaw S, Vader L, et al. antiSMASH 8.0: extended gene cluster detection capabilities and analyses of chemistry, enzymology, and regulation. Nucleic Acids Res. 2025;53(W1):W32–W38. doi:10.1093/NAR/GKAF334

33. Rodriguez-R LM, Conrad RE, Viver T, et al. An ANI gap within bacterial species that advances the definitionsdefinitionsof intra-species units. mBio. 2024;15(1). doi:10.1128/MBIO.02696-23;WGROUP:STRING:PUBLICATION

34. Yoshioka M, Akasaka N, Masuda Y, Mori M, Watanabe D. Ethanolphilic lactic acid bacterium Fructilactobacillus fructivorans as the key microorganism for fermentation of narazuke, a traditional Japanese preserved food. Appl Environ Microbiol. 2025;91(12). doi:10.1128/aem.01730-25

35. Sanghani PC, Stone CL, Ray BD, Pindel E V., Hurley TD, Bosron WF. Kinetic Mechanism of Human Glutathione-Dependent Formaldehyde Dehydrogenase. Biochemistry. 2000;39(35):10720–10729. doi:10.1021/BI9929711

36. Tragust S, Herrmann C, Häfner J, et al. Formicine ants swallow their highly acidic poison for gut microbial selection and control. Elife. 2020;9:1–25. doi:10.7554/eLife.60287

